# The cilia enriched oxysterol 7β,27-DHC is required for polycystin activation

**DOI:** 10.1101/2022.04.13.488122

**Authors:** Kotdaji Ha, Nadine Mundt, Paola Bisignano, Aide Pinedo, Erhu Cao, Jeremy F. Reiter, David R. Raleigh, Markus Delling

## Abstract

PC-1 and PC-2 form a heteromeric ion channel complex (hereafter called the Polycystin complex) that is abundantly expressed on primary cilia of renal epithelial cells. Mutations within the polycystin complex cause Autosomal Dominant Polycystic Kidney Disease (ADPKD). The Polycystin complex forms a non-selective cation channel, yet the spatial and temporal regulation of the polycystin complex within the ciliary membrane remains poorly understood, partially due to technical limitations posed by the tiny ciliary compartment. Here, we employ our novel assays to functionally reconstitute the polycystin complex in the plasma membrane. Using whole-cell and ciliary patch-clamp recordings we identified a ciliary enriched oxysterol, 7β,27-DHC, as a critical component required for activation of the polycystin complex. We identified a novel oxysterol binding pocket in PC-2 using molecular docking simulation. We also identified two amino acids within the PC-2 oxysterol binding pocket, E208 and R581, to be critical for 7β,27-DHC dependent polycystin activation in both the plasma membrane and ciliary compartment. Further, we can show that the pharmacological and genetic inhibition of oxysterol synthesis by carbenoxolone (CNX) reduces channel activity in primary cilia. Our findings identified a unique second messenger that regulates the polycystin complex. We hypothesize that cilia-enriched lipids license the polycystin complex to be functional only in the ciliary organelle, thus providing novel insights into the spatial regulation of the polycystin complex. Our results also establish a framework to target the same allosteric regulatory site in the polycystin complex to identify activators of the polycystin channels as novel therapeutic strategies for ADPKD.

---

Primary cilia are tubular structures that protrude from the surface of most mammalian cells^1^. Cilia house a specific subset of lipids and proteins designated to the cilium that must pass the transition zone at the base of the cilium, which forms a regulated diffusion barrier and provides the structural basis for the cilium as a compartmentalized organelle^2–6^. Less is known about the specific targeting of lipids to the primary cilium, but several reports suggest that the ciliary membrane harbors a specific subset of lipids (such as phosphatidylinositol-4-phosphate, PI(4)P and oxysterols), creating a specialized microenvironment for cilia specific signal transduction^2, 7^.

It is widely believed that cilia specific accumulation of proteins and lipids assigns the cilium as a unique organelle and is required to receive and transmit extracellular cues regulating diverse cellular processes ranging from early development to kidney physiology (e.g., sonic hedgehog pathway (Shh))^8–10^. Consistent with the essential role of primary cilia as cellular sensors, mutations within genes that encode ciliary proteins underlie a plethora of diseases categorized as ciliopathies affecting multiple organs and resulting in developmental, brain, motor, and kidney dysfunction^11–15^. For instance, Raleigh *et al*. identified a cilia specific oxysterol, 7β,27-DHC, required for Hedgehog (HH) signalling^16–19^. 7β,27-DHC is an oxysterol, which are cholesterol derivates with additional hydroxyl groups in the steroid ring and aliphatic chain. 7β,27-DHC is synthesized by 11b-hydroxysteroid dehydrogenase (11β-HSD)^20^, an enzyme that also catalyzes the interconversion of active cortisol and corticosterone^21, 22^. scRNA seq and proteomics study show that 11β-HSD2 is predominantly expressed along the renal tubules, suggesting that 7β,27-DHC is synthesized in the kidney^23^. However, the function of oxysterols is unknown mainly due to a lack of a tool to visualize or quantify oxysterols, but they are emerging as a physiologically diverse group of metabolites that may function as second messengers^24–26^.

PC-1 and PC-2 are the dominant ion channel expressed in primary cilia of renal epithelial cells^27–29^. PC-1 (encoded by the PKD1 gene) consists of a large extracellular N-terminal fragment (NTF) decorated with functional motifs (size~325kDa) and 11 transmembrane domains as C-terminus fragment (CFT)^30^. G-protein coupled receptor proteolytic site (GPS) cleaves PC-1 into NTF and CTF, and the intrinsic cleavage mechanism is indispensable for the function of PC-1^31–33^. PC-2 (encoded by the PKD2 gene) belongs to the transient receptor potential polycystic (TRPP) ion channel family and contains six transmembrane domains^13, 34, 35^. PC-2 is an essential subunit for the polycystin complex to function as a cation conducting ion channel^36–38^. Six transmembrane domains of PC-1 and PC-2 physically interact to form a heteromeric complex with a ratio 1:3^39^. Mutations in PKD1 and PKD2 cause autosomal dominant polycystic kidney disease (ADPKD) characterized by continued enlargement of cysts within the kidney and other organs^12, 27, 40–46^. Most ADPKD patients eventually develop end-stage renal disease (ESRD), placing a huge burden on the health care system^41, 47, 48^. While genetics and mouse models ofADPKD strongly support the idea that ciliary localization of polycystin channels is critical for channel function, the pathophysiology of the polycystin complex and primary cilia during cystogenesis is only poorly understood^8, 11^. Strikingly, restoration of PKD expression in already cystic kidneys of ADPKD animal models reverses cyst, further supporting the idea that polycystin activators are a viable path forward to develop novel therapeutics for ADPKD ^49^.

Previous studies characterizing the functional properties of PC-2 homomers and heteromers in the plasma membrane of HEK293 cells or xenopus oocytes relied on gain of function mutations in PC-2, such as PC-2_F604P_ or PC-2LN ^38, 50–52^. Further, using the PC-2 F604P mutant, we reported that the PC-1 NTF plays a pivotal role in regulating the polycystin complex^38^. However, in our initial report we failed to record from heteromeric polycystin channels containing wild type (wt) PC-2 subunits in the plasma membrane, despite reliable membrane insertion. These initial finding were surprising since wt PC-2 subunits form functional channels in primary cilia of IMCD-3 and HEK cells ^36–38, 53^. This let us hypothesize that primary cilia may contain critical co-factors required for activation of polycystin channels. We performed whole-cell and ciliary patchclamp techniques to compare the channel activity from the plasma and ciliary membrane. Using this strategy, we show that ciliary-specific oxysterol, 7β,27-DHC, binds to PC-2 to activate the polycystin complex on the plasma membrane. Furthermore, we found that pharmacologic and genetic inhibition of 7β,27-DHC synthesis reduces polycystin channel activity in primary cilia of IMCD-3 cells. This study offers compelling insights that oxysterol derivates may be further developed into PC-2 activators that can form the basis of potential ADPKD therapeutics.

## Results

### The cilia enriched oxysterol, 7β,27-DHC, activates the polycystin complex on the plasma membrane

We previously characterized the polycystin complex on the plasma membrane using a gain of function mutation, F604P, within the PC-2 subunit^38, 51, 52^. In order to identify potential activators of the polycystin complex, we developed stable cell lines that co-express WT PC-2 together with sPC-1, in which the endogenous signal peptide has been replaced by the Ig k-chain secretion sequence and HA tag ^38, 51, 54^ (Figure 1A). Live cell staining with an anti HA antibody revealed that the heteromeric complex traffics to the plasma membrane of HEK293 cells (Figure 1B).

**Figure 1.**
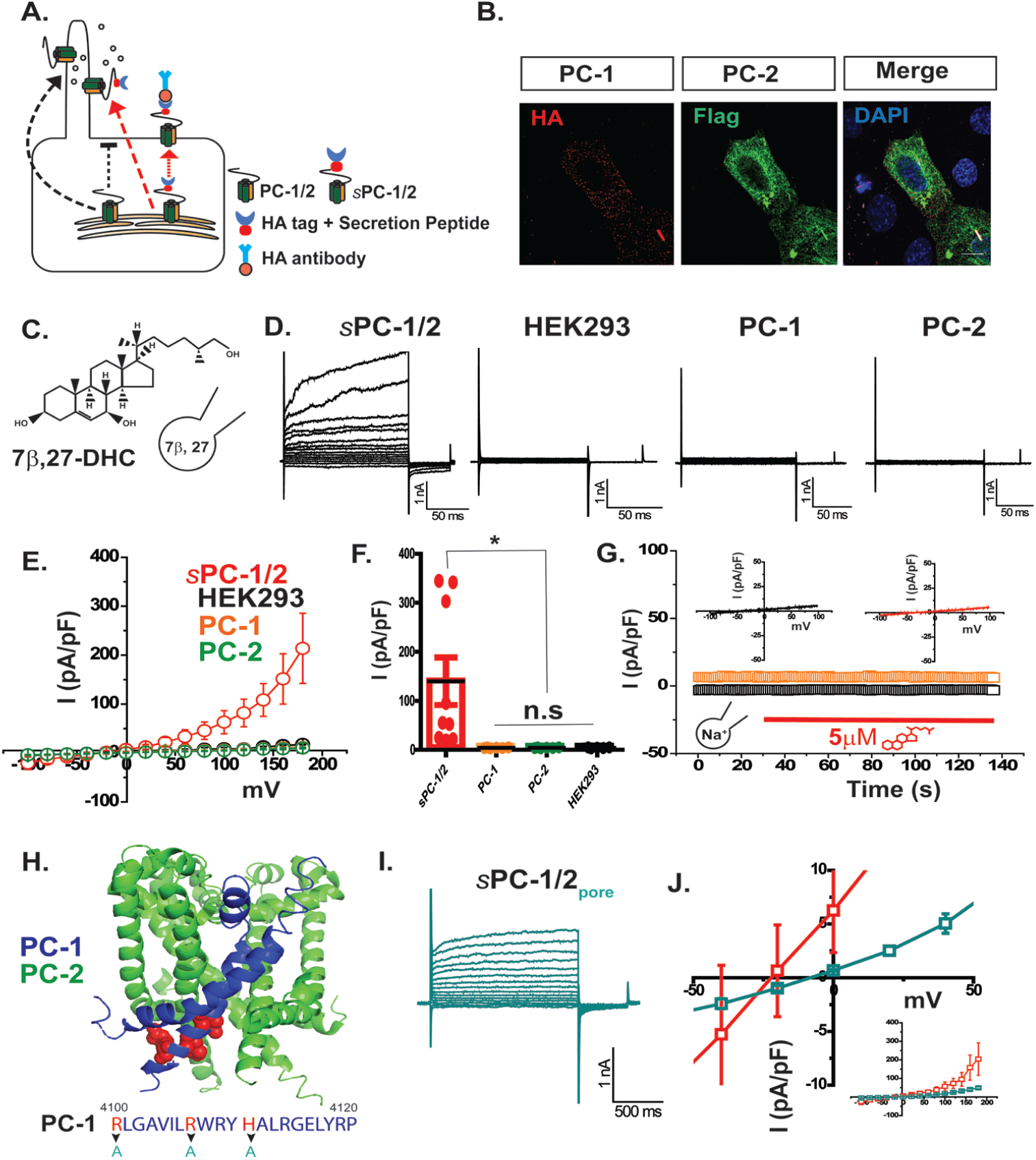
ciliary oxysterol, 7β,27-DHC, activates the polycystin complex on the plasma membrane. A. Schematic illustration of strategy to target the polycystic complex on the plasma membrane. Black dotted arrow indicates the endogenous trafficking system of PC-1 and PC-2. Red dotted arrow shows the novel trafficking system we developed previously using secretion 38. We added secretion peptide in the N-terminus of PC-1 to target the surface membrane and HA tag for the detection. B. Immunofluorescent image of polycystin complex on the surface membrane. HA-tagged PC-1 (HA-antibody:red), Flag-tagged PC-2 (Flag-antibody:green) and Nucleus (DAPI:blue). Images are merged to show colocalization. C. Molecular structure of ciliary oxysterol, 7b,27-DHC. We added 5µM 7b,27-DHC in the glass electrode to inject intracellularly for the whole-cell patch clamp. D. Whole-cell patch clamp traces recorded from transiently expressed sPC-1/2, PC-1, PC-2 cells and HEK 293 cells. The voltage step pulse from −100 mV to +180 mV in +20 mV increments was applied for the whole-cell patch clamp with 0 mV holding potential. The step pulse was given for 150 ms at each step pulse. The tail pulse was given at −80 mV after each voltage step pulse injection during 50 ms. E. Current (I)-voltage (V) relationship for sPC-1/2, PC-1, PC-2, and HEK 293. Each value was obtained at the maximum currents (pA) from −100 mV to +180 mV voltage step pulse and divided by capacitance (pF); sPC-1/2 (red, n=8), PC-1 (orange, n=12), PC-2 (green, n=10), and HEK293 (n=15). F. Comparison of the current density at +180 mV from Figure 1E. G. Extracellular treatment of 7β,27-DHC. 5µM 7β,27-DHC was added in the extracellular solution and perfused onto the cell membrane. After applying, sPC-1/2 overexpressed HEK293 cells were recorded using whole-cell patch clamp. The ramp pulse was applied from −100 mV to +100 mV during 500 ms. Each value (pA) obtained at −100 mV (black blank square) and +100 mV (orange blank square) divided by capacitance (pF) was picked and plotted over time. After 30 seconds recording using ramp pulse, 5µM 7β,27-DHC was perfused for 70 seconds. Small graphs are the I-V relations before (left) and after (right) application. H. Cryo-EM structure of PC-1 and PC-2 heteromeric channel (PDB: 6A70) and the pore region of PC-1 sequence alignment. We substituted R4107, H4111, and R4104 with alanine. The location of R4107, H4111, and R4104 is shown in sphere (red). I. Whole-cell patch clamp trace recorded from sPC-1/2 pore mutant-overexpressing HEK293 cell. The voltage step pulse is identical to other tests. J. Comparison of current (I)-voltage (V) relationships for sPC-1/2 pore mutant (green) and sPC-1/2 (red). We zoomed in I-V relationship from −50 mV to +50 mV. The whole I-V curve is shown below. All summary data, mean ±SEM.

After confirming plasma membrane insertion of sPC-1/2, we tested a variety of cilia enriched lipids, including oxysterol isoforms, in their ability to activate the polycystin complex. We performed whole-cell patch-clamp recordings and applied the lipids in the bath solution or patch pipette solution, allowing us to selectively apply the lipid to either the extracellular or intracellular side of the polycystin complex (**Suppl Data 1**). Our initial screen of 10 cilia enriched lipids identified 7β,27-DHC as the only potential activator (**Suppl Data 1A-C**). When we included 5µM 7β,27-DHC in the intracellular solution of the patch pipette, we recorded an outward rectifying current from sPC-1/2 expressing HEK293 cells (139.9±48.3 pA/pF, n=9) (**Figure 1D-F**). In contrast, untransfected HEK293 cells did not show any currents with the intracellular 5µM 7β,27-DHC (9.3 ±1.3 pA/pF, n=15), suggesting that 7β,27-DHC specifically activates the polycystin complex (Figure 1D-F).

To test whether 7β,27-DHC can activate either PC-1 or PC-2 alone in our heterologous expression system, we performed patch-clamp recordings on HEK293 cells either expressing sPC-1 or PC-2 with 5µM 7β 27-DHC in the patch pipette. PC-1 or PC-2 expressing HEK cells failed to generate currents above background (PC-1, 8.3 ± 2.3 pA/pF, n=10; PC-2, 10.1 ± 0.7 pA/pF, n=10) (Figure 1D-F). These findings indicate that both PC-1 and PC-2 are required to form a functional channel in the plasma membrane^38^.

To characterize the mechanism by which 7β,27-DHC activates the channel complex, we acutely applied 5µM 7β,27-DHC to the extracellular bath solution of sPC-1/2 expressing HEK cells and performed whole-cell recordings using a ramp pulse. However, extracellular application of 7β,27-DHC did not activate sPC-1/2 in HEK293 cells (10.4 ± 2.3 pA/pF, n=24) (**Figure 1G**). Collectively, these findings suggest that the 7β,27-DHC acts via the cytoplasmic leaflet to modulate channel activation.

To confirm the behavior of the polycystin complex, we introduced three mutants, R4100A, R4107A, and H4111A of PC-1 (sPC-1/2_pore_) and recorded sPC-1/2_pore_ over-expressing HEK293 cells using the whole-cell patch-clamp with intracellular 5µM 7β,27-DHC. R4100, R4107, and H4111 in the S6 of the PC-1 transmembrane domain suggested interfering with cation permeation^38, 39^. sPC-1/2 pore resulted in outward currents with shifted reversal potential, suggesting that introducing the mutants modified the characteristics of the polycystin complex (**Figure 1I-J**). We conclude that 7β,27-DHC activates wild type polycystin and overcomes the dependency on the F604P gain of function mutation for plasma membrane recordings.

### 7β,27-DHC activates the polycystin complex in a dose-dependent manner

We next hypothesized that 7β,27-DHC may interact directly with the polycystin complex. We measured polycystin activation in the inside-out single-channel configuration in the sPC-1/2 over-expressing HEK293 cell membrane and applied 1µM, 2.5µM, 5µM, and 50µM 7β,27-DHC to the bath solution (Fig2A). In this configuration the cytoplasmic leaflet is exposed to the bath solution. In single-channel recordings, 7β,27-DHC activated sPC-1/2 in a dose-response manner (Figure 2B. G_1μM_ 57.2 ± 1.9 pS, n=5; G_2.5 μM_ 66.4 ± 1.0 pS n=5; G_5μM_ 63.75 ±9.65 pS n=5; G_50μM_ 76.87 ± 4.28 pS n=5). Consistently, a higher concentration 7β,27-DHC elicited a dose-dependent increase in the open probability, characterized by quick transitions between open and close states (**Figure 2C**). Of note, 50μM 7β,27-DHC also increased open probability at −100 mV membrane potential. Thus, 7β,27-DHC behaves as a gating modifier to promote the open state and conductance of the polycystin complex. We thus demonstrated for the first time that cilia-specific oxysterols activate the polycystin complex.

**Figure 2.**
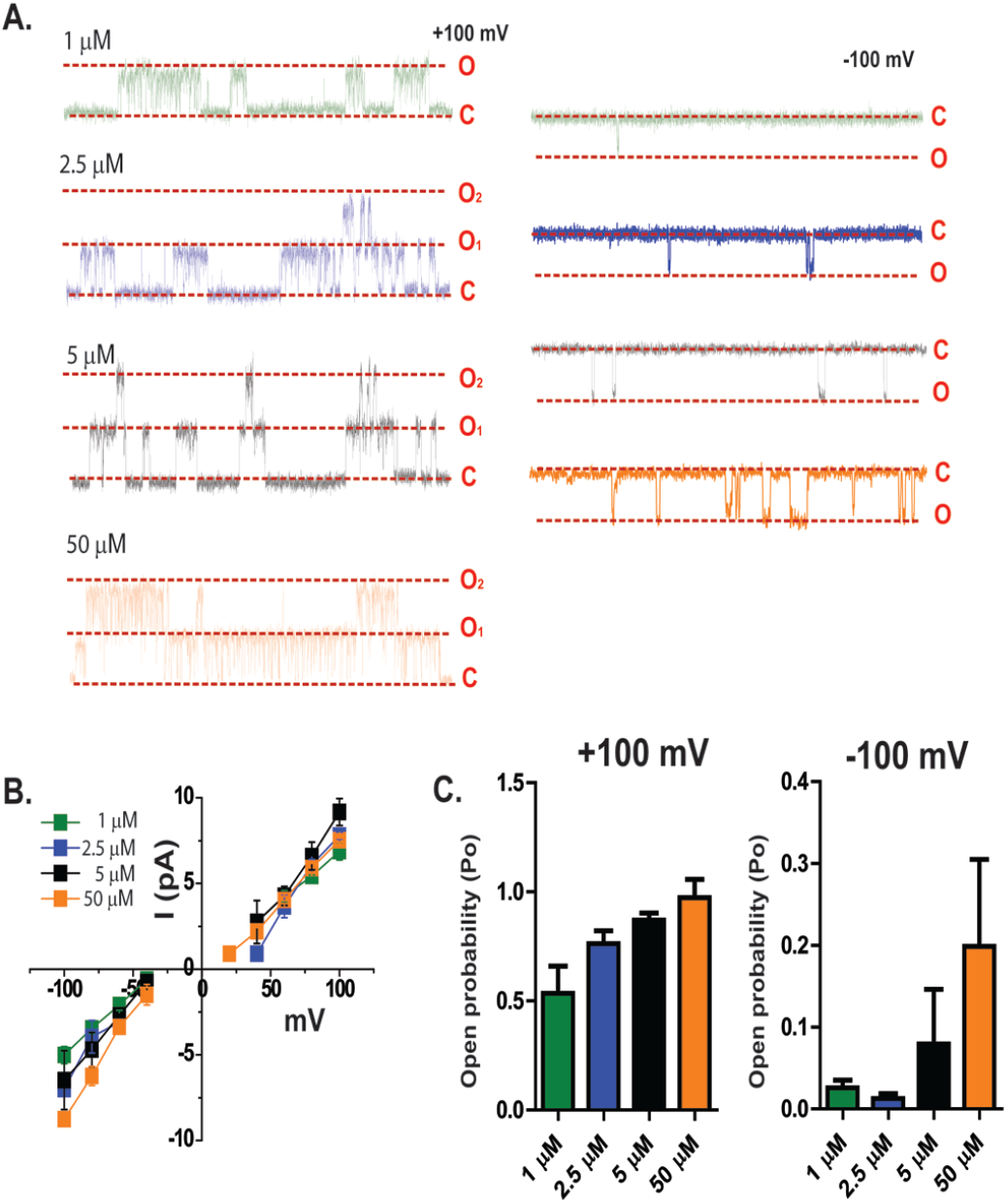
Dose dependent activation of sPC-1/2 by 7β,27-DHC. A. Representative inside-out single channel recordings of sPC-1/2 overexpressing HEK293 cells with 1µM, 2.5 µM, 5 µM, and 50 µM 7β,27-DHC intracellular application. Red dotted lines indicate close (C) and open (O) state of single channel. B. Single channel conductance of with 1µM, 2.5 µM, 5 µM, and 50 µM 7β,27-DHC intracellular application. n=5-6. C. Normalized absolute open probability of sPC-1/2 obtained from −100 mV to +100 mV during 10s recording. The absolute open probability was normalized to the maximum open probability. n=5-6.All summary data, mean ±SEM.

### Ciliary oxysterol, 7β,27-DHC, further potentiates ciliary polycystin channels

The polycystin complex shows basal activity in the ciliary membrane^37, 38^, raising the question whether exogenously applied 7β,27-DHC can further potentiate the activity of the polycystin complex in the ciliary membrane. To test this hypothesis, we performed excised ciliary inside-out patch-clamp recordings (**Figure 3B**) to measure activation of endogenous polycystin channels by 7β,27-DHC. Ciliary membrane was excised from ARL13B-EGFP expressing mIMCD-3 cells and the intraciliary leaflet was perfused with 7β,27-DHC. We found that in the presence of 7β,27-DHC ciliary polycystin channels already started to open at +40 mV and −20 mV, while channels remain closed at these potentials without exogenous addition of 7β,27-DHC. This strongly suggests that 7β,27-DHC further potentiates the ciliary polycystin complex (**Figure 3A**). Furthermore, the absolute open probability of ciliary polycystin channel with 7β,27-DHC was right-shifted at negative potential, indicating channel activation in the physiological range (**Figure 4C**). The conductance of ciliary polycystin channels did not change with 7β,27-DHC (G_cilia+7β,27-DHC_: 86± 12.0 pS, n=5; G_cilia_ :89.0 ± 5.6 pS, n=6). Taken together, these findings demonstrate that 7β,27-DHC can further potentiate ciliary polycystin channels at physiological membrane potentials.

**Figure 3.**
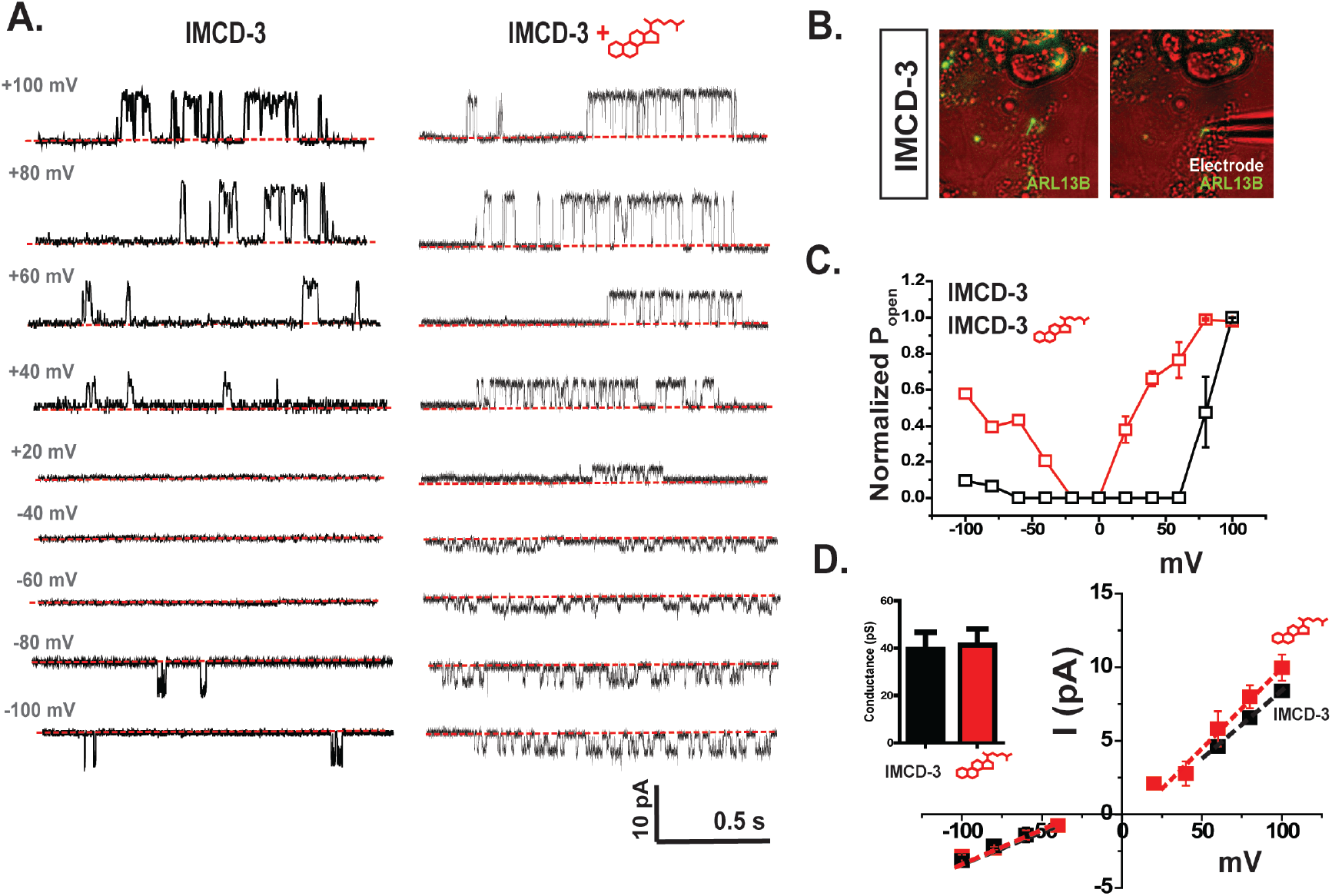
Ciliary oxysterol, 7β,27-DHC, also activates ciliary currents. A. Representative excised ciliary inside-out single channel recordings of primary cilia in IMCD-3 (left) and 5µM 7β,27-DHC perfused recordings (right) Red dotted line indicates close state of channel. B. Images of ARL13B-GFP expressing primary cilium in IMCD-3 cell (left) and patched primary cilium with a glass electrode (right). C. Normalized absolute open probability of ciliary channels from IMCD-3 cells. D. Single channel conductance of the ciliary channel recording shown in Figure 3A. Black and red squares indicate the averaged currents amplitudes from ciliary channel from IMCD-3 cells (n=6) and 5µM 7β,27-DHC perfused ciliary channel (n=5), respectively. Dotted line indicates the fitting to the linear equation. Small bar graph on left shows the comparison of conductance comparing primary cilia to 5µM 7β,27-DHC treated primary cilia. All summary data, mean ±SEM.

**Figure 4.**
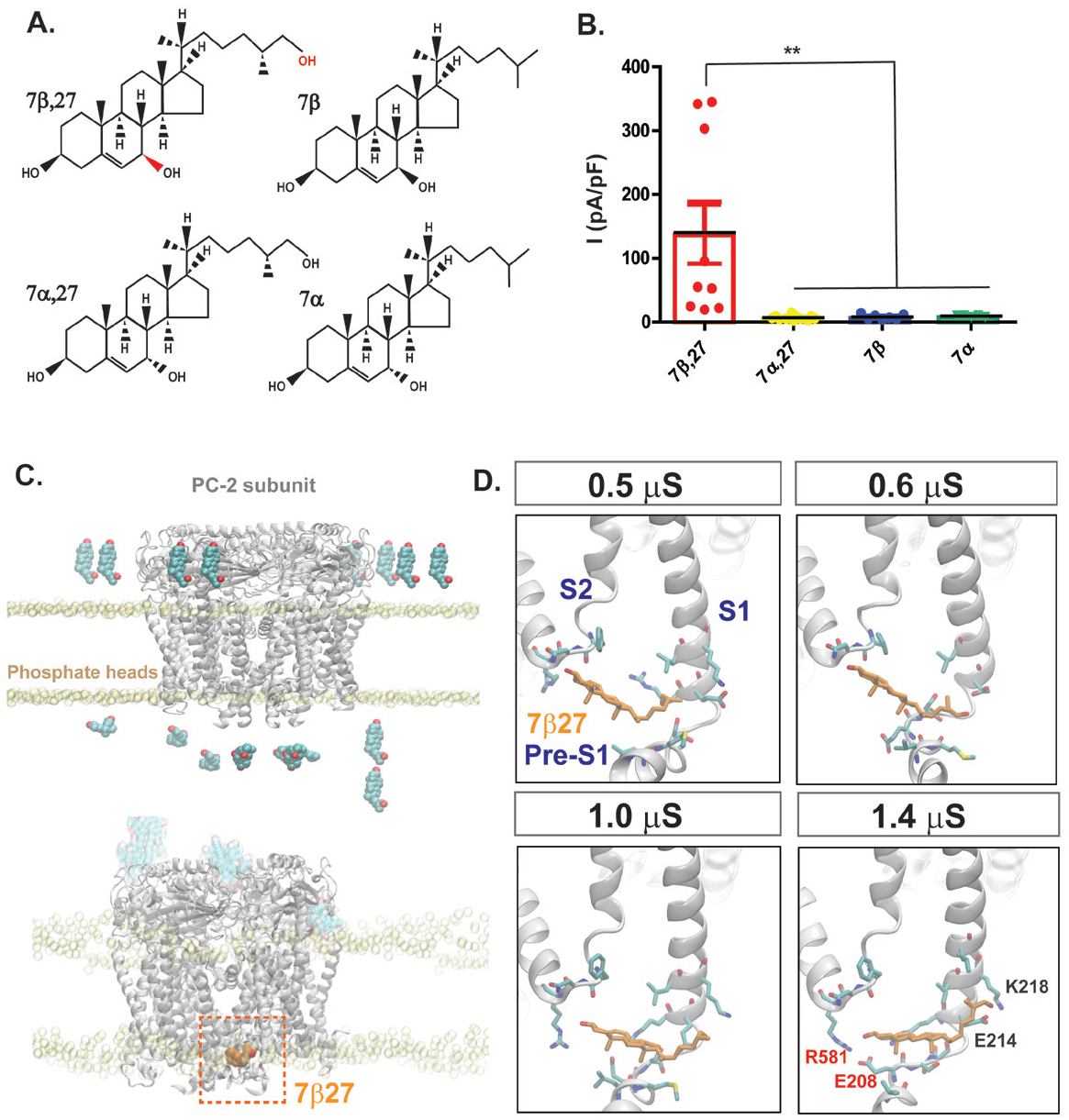
7β,27-DHC binds to PC-2 subunit. A. Structural similarity of different oxysterols. Molecular structures of three oxysterols, 7α, 7β,27-DHC, and 7β based on the structural similarity to 7β,27-DHC. Hydroxyl groups colored in red indicate the distinguished structure in 7β,27. B. Current density at +180 mV from the whole-cell recordings of HEK293 transiently expressing sPC-1/2. 5µM 7α, 7β,27-DHC, and 7β were injected to HEK293 cells transiently expressing sPC-1/2 intracellularly. C. Computed oxysterol binding simulation based on PC-2 structure (PDB: 5T4D). Top image shows the arrangement of eight 7β,27-DHC molecules from extracellular and intracellular side for simulation and bottom image shows the potential docking of one 7β,27-DHC molecule (orange) to the intracellular loops and portions of PC-2. D. 7β,27-DHC binding to PC-2 over time.

### 7β,27-DHC specifically binds to the PC-2 subunit

Although 7β,27-DHC activates both heterologously expressed sPC-1/2 and ciliary polycystin channels, it remains unclear whether activation is mediated through a direct interaction with the channel complex. To ask whether oxysterols with a similar molecular structure to 7β,27-DHC can activate sPC-1/2, we applied 5µM 7β hydroxycholesterol (7β-HC), 7α-HC, and 7α,27-DHC (**Figure 4A**) to the intracellular solution in whole-cell patch-clamp recordings of sPC-1/2 overexpressing HEK293 cells. Interestingly, 7β-HC, 7α, and 7α,27-DHC failed to generate significant outward rectifying currents compared to 7β,27-DHC (Figure 4B), thereby suggesting that the specific location and orientation of the hydroxyl groups is critical to activate the polycystin complex.

To assess whether 7β,27-DHC may directly bind to the polycystin complex, we performed molecular docking simulation using 7β,27-DHC and the published PC-2 structure35. This approach is supported by a recent study which identified PC-2 as a cholesterol-binding protein55. We focused on the cytoplasmic domains of the PC-2 complex because extracellular application of 7β,27-DHC did not potentiate the polycystin complex (**Figure 1G**). As shown in Figure 4C, D the molecular docking simulations identified a putative oxysterol binding pocket formed by the pre-S1 helix and S4 and S5 linker. In particular, E208 within pre-S1 and R581 in the S4-S5 loop appeared as critical amino acids to interact with 7β,27-DHC (**Figure 4D**).

To test the importance of E208 and R581 for 7β,27-DHC-dependent channel activation, we mutated both amino acids into alanine (hereafter called sPC-1/2_E208A_ and sPC-1/2_R581A_). We confirmed that both mutants successfully localize in the plasma membrane of sPC-1/2_E208A_ and sPC-1/2_R581A_ stably expressing HEK293 cells (**Figure 5A, B**). Despite membrane expression, sPC-1/2_E208A_ (8.7 ± 0.8 pA/pF, n=10) and sPC-1/2_R581A_ (6.8 ± 0.9 pA/pF, n=12) failed to generate any currents above background in the presence of 5µM 7β,27-DHC, strongly suggesting that these two residues are critical for 7β,27-DHC activation (**Figure 5C top and D**). To further confirm that mutation of E208 or R581 does not impair general functionality of the channel complex, we additionally introduced the gain of function mutation F604P (sPC-1/2_E208A-F604P_ and sPC-1/2_R581A-F604P_) (**Figure 5C, bottom and D**). Albeit smaller than PC-2_F604_, both double mutants still produced an outwardly rectifying current in whole cell recordings of sPC-1/2_E208A-F604P_ (33.0 ± 10.0 pA/pF, n=6) and sPC-1/2_R581A-F604P_ (44.6 ± 8.9 pA/pF, n=5) suggesting that E208 and R581 are specifically required for 7β, 27-DHC dependent activation.

**Figure 5.**
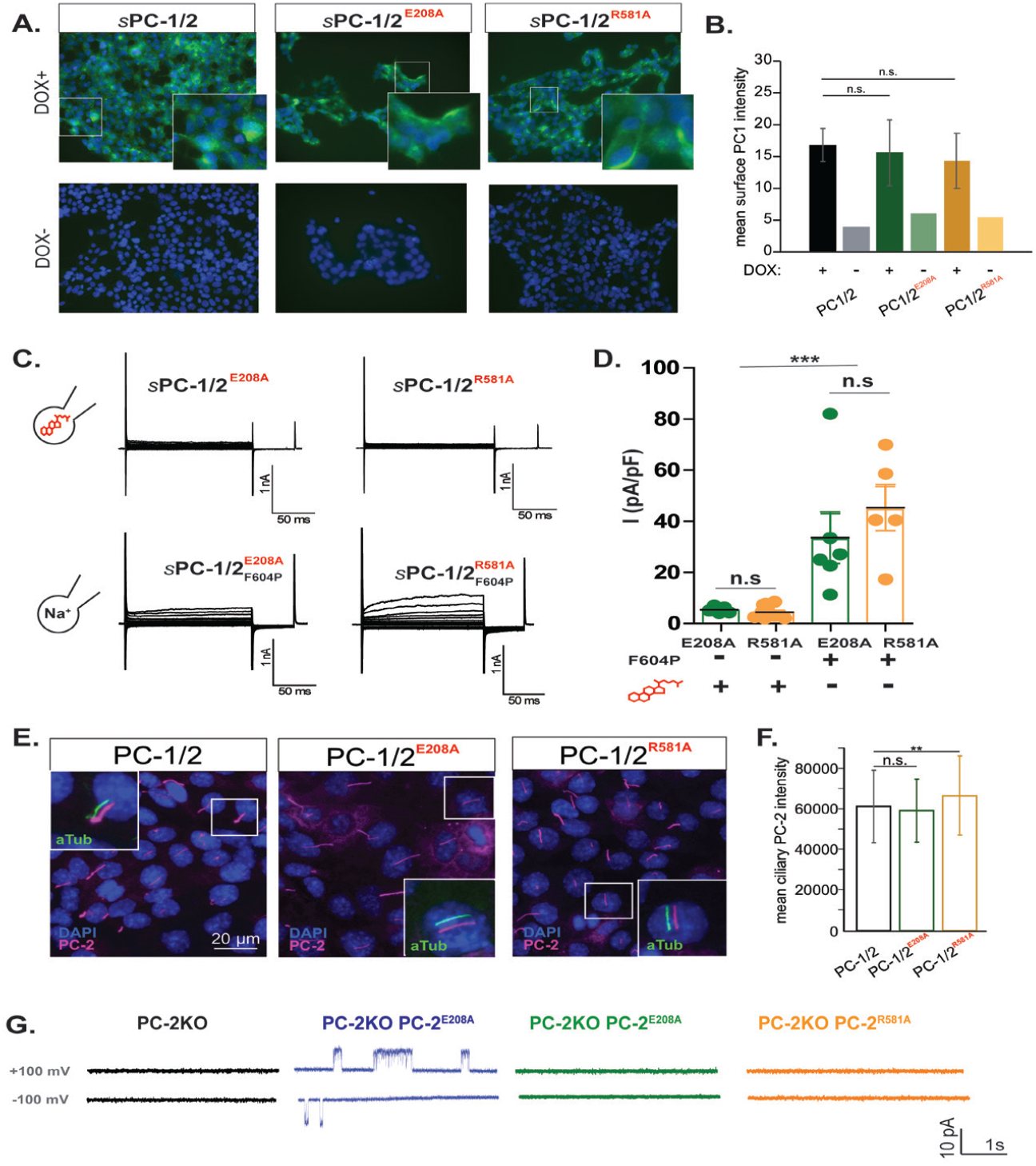
E208 and R581 are the potential binding sites for 7β,27-DHC to PC-2. A-B. Surface HA staining in stable HEK cell line and imaging analysis. We used PC-1 as an indicator for the successful PC-2 trafficking. C. Whole-cell patch clamp recordings from two oxysterol binding mutants, E208A and R581A and double mutant with gain of function mutation, F604P. The same voltage step pulse protocol was applied to test the binding mutants for the recordings. 5µM 7β,27-DHC was added in the intracellular solution for the binding mutants test (sPC-1/2_E208A_ and sPC-1/2_R581A_). 7β,27-DHC was excluded in the intracellular solution for the binding mutants with F604P test (sPC-1/2_E208A-F604P_ and sPC-1/2_R581A-F604P_). D. Comparison of current density at +180 mV from the whole cell patch clamp recordings of two oxysterol binding mutant test. The current amplitudes at +180 mV from two oxysterol binding mutants, E208A (green, n=10) and R581A (yellow, n=12) were compared to the double mutants, E208A-F604P (green, n=6) and R581A-F604P (yellow, n=5). C. Ciliary expression of PC-2_E208A_ and PC-2_R581A_. Two binding mutants were overexpressed in primary cilia of IMCD-3 cells. E-F. Immunofluorescent imaging analysis of ciliary expression of PC-2_E208A_ and PC-2_R581A_. E. Representative excised ciliary single channel recordings of PC-2 KO IMCD-3 cells and PC-2, PC-2_E208A_ and PC-2_R581A_ overexpressing PC-2 KO IMCD-3 cells.

To test whether sPC-1/2_E208A_ and sPC-1/2_R581A_ can still elicit ciliary currents, we performed excised ciliary single-channel recordings from IMCD3 cells expressing either mutant. Both mutants still localize on the ciliary membrane (**Figure 5E, F**). To avoid recordings from endogenous polycystin channels, we performed the ciliary patch-clamp recordings using IMCD-3 cells with ablated PC-2 expression^36, 38^. We reintroduced wt PC-2, PC-2 E208A and PC-2_R581A_ (**Figure 5G**, PC-2 n=4; PC-2_E208A_ n=21; PC-2_R581A_ n=22). As shown in **Figure 5G**, re-expressed ciliary PC-2 shows basal activity in the ciliary membrane, presumably elicited by endogenous 7β,27-DHC. In contrast, both E208 and R581 mutants fail to produce basal activity. These results further support the idea that E208 and R581 are critical within the 7β,27-DHC binding pocket.

### PC-2 is the only oxysterol-sensitive PC-2 family protein

To test whether PC-2 is the only oxysterol-sensitive protein among the PC-2 family, we tested activation of the close homologue PC-2L1 with 7β,27-DHC. Based on protein sequence alignments, E208 and R581 in PC-2 are not conserved in PC-2L1 and PC-2L2 (**Figure 6A**), thereby indicating that PC-2 is the only 7β,27-DHC sensitive polycystin member. We performed inside-out single-channel recordings in HEK293 cells transiently expressing PC2-L1-EGFP. Notably, PC-2L1 did not show further potentiation with 5µM 7β,27-DHC (**Figure 6B**). In addition, PC-2L1 did not result in significantly different conductance in outward (G_PC-2L1+7β,27-DHC_: 208.0 ± 7.8 pS, n=6, G_PC-2L1_; 205.8 ± 10.8pS, n=5) and inward currents (G_PC-2L1+7β,27-DHC_: 89.0± 22.0 pS,n=6, G_PC-2L1_; 107.0 ± 12.0 pS, n=5) (**Figure 6C, D**). Furthermore, the absolute open probability of PC-2L1 was not affected by 7β,27-DHC (**Figure 6E**). Thus, our results indicate that 7β,27-DHC is specific for PC-2.

**Figure 6.**
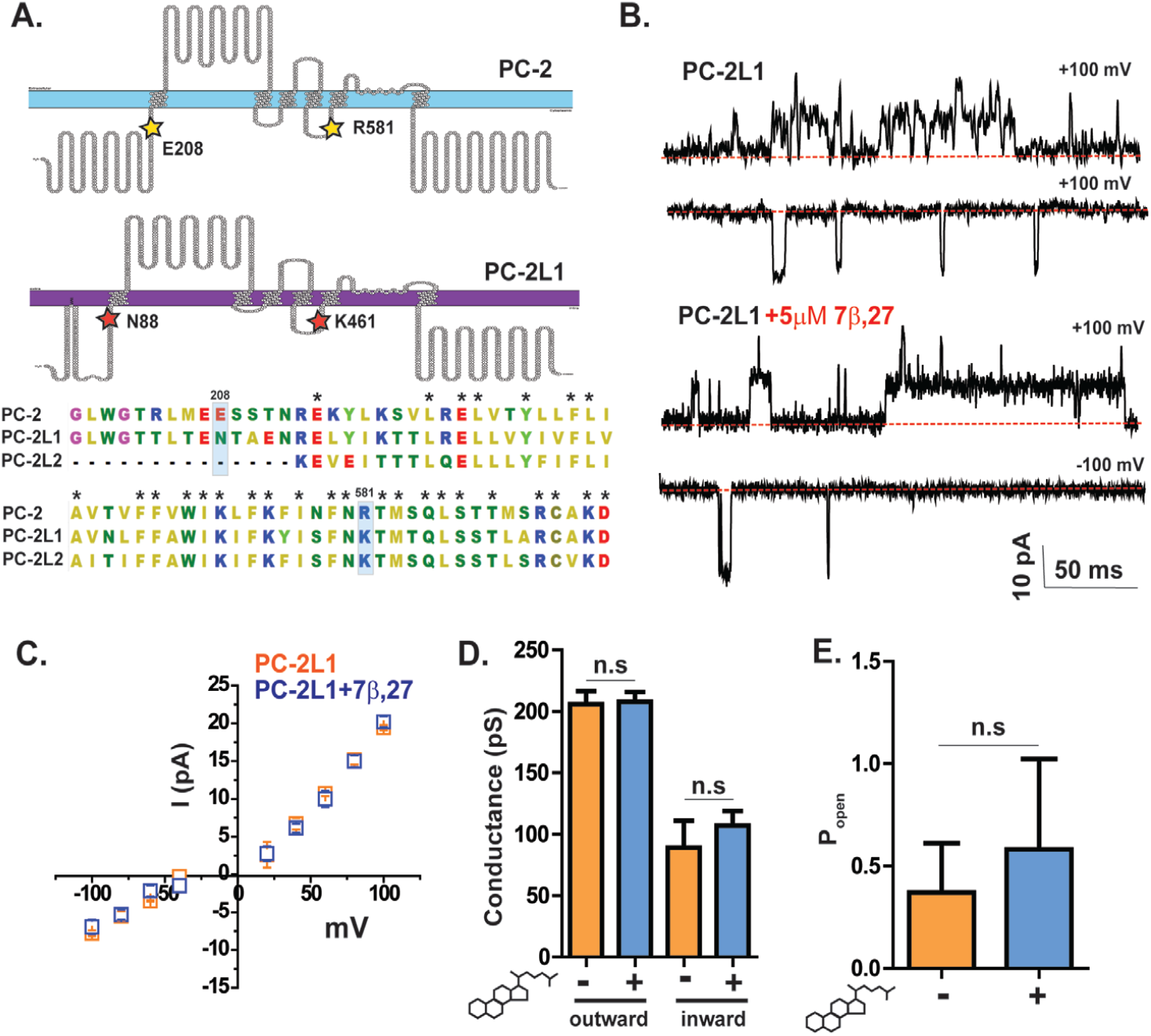
PC-2 is the only oxysterol-sensitive PC-2 family protein. A. Schematic diagram of PC-2 and PC-2L1 using protter protein program and protein alignment comparison of PC-2 family proteins, PC-2, PC-2L1 and PC-2L2. Yellow stars indicate the location of oxysterol binding mutants, E208 and R581 on the topology of PC-2. Red stars indicate N88 and K491, the location of corresponding oxysterol biding sites in PC-2L1. Black star on the protein sequences indicates 100% conserved amino acids among PC-2 family proteins. B. Representative single channel recordings of PC-2L1 at −100 mV and +100 mV with (bottom) and without (top) intracellular 5µM 7β,27-DHC Red dotted line indicates the closed channel state during recordings. C. The current amplitudes collected from −100 mV to + 100 mV. Data are fitted to the linear equation to calculate conductance (γ). Blue and orange dotted lines indicate the linear fitting to the average current amplitudes of PC-2L1 with (blue, n=6) or without (orange, n=5) intracellular 5µM 7β,27-DHC. D. Comparison of PC-2L1 conductance obtained from inward and outward currents with (blue, n=6) or without (orange, n=5). E. Comparison of PC-2L1 open probability at −100 mV with (blue, n=6) and without (orange, n=5) the intracellular 5µM 7β,27-DHC.

### Synthesis of ciliary oxysterols by HSD11B enzymatic activity is essential for polycystin activation

Having established that 7β,27-DHC is a crucial component to activate the polycystin complex in the plasma membrane and the ciliary membrane, we next asked whether 7β,27-DHC is required for polycystin activity in primary cilia. 11b-Hydroxysteroid dehydrogenase (HSD11β) is the oxysterol synthase essential for generating ciliary oxysterols^17, 56^. Carbenoxolone (CNX), a component of licorice, is known to inhibit the activity of 11β-hydroxysteroid dehydrogenase^17^ (**Figure 7A**). A recent study demonstrated that inhibition of β2-HSD11 by 400 nM CNX impaired activation of the hedgehog pathway in primary cilia^17^. We thus hypothesized that CNX could also impair ciliary polycystin activity by lowering 7β,27-DHC levels. As shown in **Figure 7B**, incubation of IMCD3 cells with CNX for 72 hrs completely abrogated endogenous polycystin channel activity (**Figure 7B**), even during long recordings (n=20). Next, we assessed the effects of genetic inhibition of 7β,27-DHC by applying small interfering RNA (siRNA) to knock down endogenous expression of β2-HSD11. We also performed the double knockdown of β1-HSD11 and β2-HSD11 to avoid a potential compensation mechanism for β2-HSD11. Genetical depletion of both enzymes resulted in complete loss of ciliary polycystin currents compared to the scrambled siRNA (**Figure 7D, E**). These findings strongly demonstrate that oxysterol synthesis is required for ciliary polycystin activity in IMCD-3 cells.

**Figure 7.**
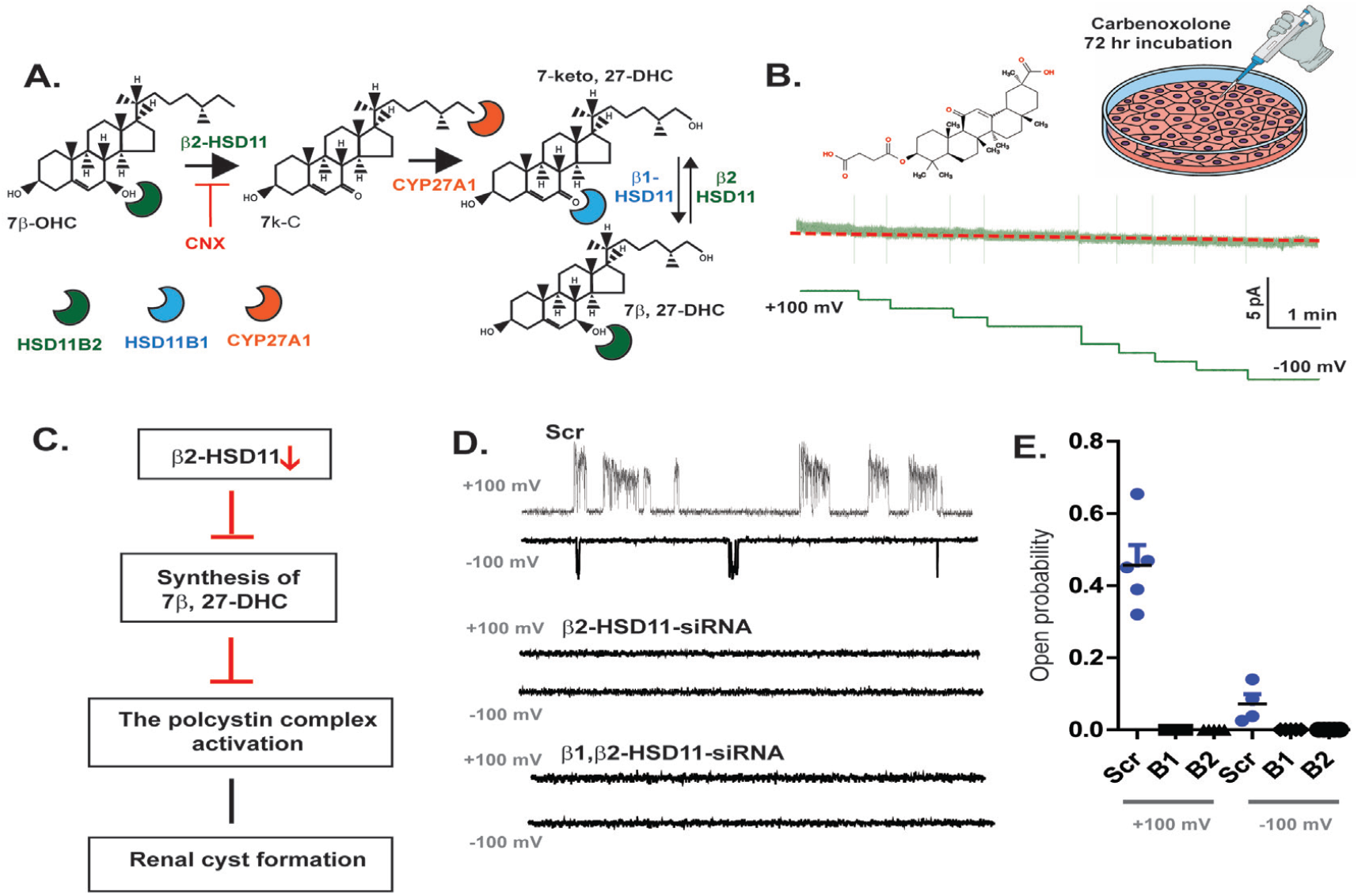
Synthesis of 7β,27-DHC is crucial to activate ciliary currents. Schematic pathway of 7β,27-DHC synthesis. Schematic diagram illustrates enzymatic activity of 11β-Hydroxysteroid dehydrogenase type 1 (β1-HSD11) to convert 7-keto, 27-DHC to 7β,27-DHC. Carbenoloxone (CNX) inhibits enzymatic activity of β2-HSD11. Excised ciliary single channel recording of IMCD-3 cells after 72-hour Carbenoloxone (CNX) incubation. Holding potentials were given from +100 mV to −100 mV for 13 min. Red dotted line indicates the close state of channel. C. Schematic diagram of inhibition mechanism for the polycystin complex activation. D. Excised ciliary single channel recordings from scrambled siRNA control primary cilium (top), β1-HSD1-siRNA treated primary cilium (middle), and β2-HSD11-siRNA treated primary cilium (bottom). E. Measurement of open probability of ciliary recordings at +100 mV and −100 mV potentials after HSD11siRNA transfection. We counted open event of the single channel recordings at +100 mV and −100 mV. Scr n=5, β1-HSD11 n=21, β2-HSD11 n=23.

## Discussion

Here we identified for the first time a cilia-specific oxysterol, 7β,27-DHC, as a critical cofactor for polycystin activation. Our study was initiated by the puzzling observation that wt PC-2 does not produce any functional channels in the plasma membrane when co-expressed with sPC-1, despite efficiently trafficked to the plasma membrane^38^. This made us speculate that cilia-enriched lipids may be required for channel activation. Our current model is that cilia-specific lipids confer spatial specificity for activating the polycystin complex only in the ciliary membrane. While PC-2 has been reported in several subcellular locations, including the ER^57^, the further dependence on organelle-specific cofactors licenses channel activation only in the cilium. This is a common concept in channel physiology: for instance, the mucolipins or TRPML’s, the closest TRP channel homologues to PC-2, require PI(3,5)P2 for activation, which is exclusively found in the endosomal membrane, thus restricting TRPML activity to the endosome^58–60^. While phosphatidylinositol phosphates are well established as second messengers required for TRP channel activation, the cilia enriched PI(4)P did not activate the polycystin complex (**Suppl Figure 1**). Our results for the first time explore oxysterols as critical factors for ion channel physiology. It is currently unclear whether oxysterols are constitutive components of the ciliary membrane or can function as second messengers which are dynamically regulated on short time scales. Novel biosensors are needed to study the regulation of oxysterols in subcellular compartments.

This study suggests that cilia-enriched oxysterol is a critical cofactor to activate the ciliary polycystin complex. While other oxysterols structurally related to 7β,27-DHC failed to activate the polycystin complex, we cannot rule out that other cofactors can contribute to channel activation. Using molecular docking simulation, we predicted a potential binding site of 7β,27-DHC in PC-2. Interestingly Wang *et al*., also predicted a location near E208 and R581 as a potential phospholipid or cholesterol-binding pocket in the human PC-2^61^. E208 and R581 are not conserved among other PC-2 family proteins, including PC-2L1 and PC-2L2, suggesting that PC-2 is a unique a lipid-binding protein. Indeed, a proteomics study identified PC-2 as the only sterol-binding protein among the PC-2 family proteins^55^.

However, the clinical correlation between oxysterols, polycystins, and ADPKD remains unclear. Although pharmacologic or genetic inhibition of synthesis of 7β,27-DHC resulted in a loss of ciliary currents, β2-HDS11 knock out mice do not demonstrate cystogenesis in the kidney^62^. 7β,27-DHC synthesizing enzyme β2-HSD11 is highly expressed in the developing mammalian kidney^22^ and serves numerous physiological functions^21, 22, 63^. Notably, the inhibition of glucocorticoid synthesis by β2-HSD11 expression is well explored, and deficiency of β2-HSD11 can cause inappropriate glucocorticoid activation of renal mineralocorticoid receptors (MR)^64, 65^. Clinically, the deficiency of β2-HSD11 in the kidney causes Apparent Mineralocorticoid Excess (AME) syndrome, which is characterized by hypertension and hypokalemia^20, 66–69^. Moudgil *et al*., reported renal cysts in an AME patient, showing that inhibited β2-HSD11 enzymatic activity may lead to renal cyst formation^70^. Furthermore, a high dose of glucocorticoid in newborn mice induced polycystic kidney disease^71^. Thus, the clinical evidence linking inhibition of 7β,27-DHC and renal cyst formation still remains unclear.

This study shows that cilia-specific oxysterol, 7β,27-DHC, can potentiate polycystin channel activity. We also show that 7β,27-DHC binding to PC-2 is enantioselective, highlighting the specificity of the interaction. Recent research demonstrated that re-expression of PC-2 can reverse renal cystogenesis, showing that the activity of polycystins suppresses cilia-dependent cyst-activating (CDCA) signals^49^. Our study also suggests that developing PC-2 activators specifically targeting the oxysterol binding site is a promising therapeutic approach to compensate for ADPKD-causing loss of function mutations in PC-2.

## Methods

### Cell culture and transfection

Tetracycline-inducible human embryonic kidney (HEK) 293 cells and mouse inner medullary collecting duct 3 (mIMCD-3) were used for whole cell patch clamp and ciliary excised single channel recordings, respectively. HEK293 were cultured in Dulbecco’s Modified Eagle Medium (DMEM; Gibco) supplemented with 10% fetal bovine serum (FBS;) and 0.1% penicillin-streptomycin at 37C in a CO2 incubator (Heraeus). mIMCD-3 were cultured in F-12K Nutrient Mixture (1X) Kaighn’s Modification (Gibco) supplemented with 10% FBS and 0.1% penicillin-streptomycin. For ciliary induction, mIMCD-3 cells were incubated in OPTI-MEM for 2 days in CO2 incubator. sPC-1/2 construct was cloned into pTRE vectors and transfected using lipofectamine 2000 into HEK293 cells, according to the manufacturer’s instructions. sPC-1/2 transfected HEK293 cells were treated with 1 uM of doxycycline 12 hours before the experiment. For the patch clamp experiment, the cells were seeded into the path clamp chamber. For the ciliary patch clamp test, mIMCD-3 cells were transfected with ARL13B-GFP using lipofectamine LTX, according to the manufacturer’s instructions.

### Immunocytochemistry and imaging

All procedures were performed at room temperature. Cells were fixed in 3.2 % paraformaldehyde for 10 min, permeabilized in 0.2 % Triton X-100 for 10 min, and blocked-in blocking solution (BS) that contains 5% FCS, 2% BSA, 0.2% Fish Gelatin, 0.05% NaN3 in PBS) for 30 min. Cells were then incubated with primary antibody solution (BS plus primary antibody at 1:1000 dilution) for 1 hour. Cells were washed two times with PBS-/-before incubation with secondary antibody solution (BS plus secondary antibody and Hoechst at 1:1000 dilution) for 1 hour. Cells were repeatedly washed with PBS-/- and mounted onto glass coverslips for imaging.

To stain for surface HA-tag expression, nonfixed cells were incubated in rat anti-HA (3F10, Roche) primary antibody at a concentration of 1:100 in Leibovitz’s L-15 medium for 20 min. After washing with antibody-free L-15 medium the staining protocol preceded as described above.

Mounted cells were imaged on a Nikon Ti inverted fluorescence microscope with CSU-22 spinning disk confocal and EMCCD camera. Z-stacks were acquired to visualize the full z-dimension of each cilium. To compare trafficking between wildtype and mutant PC2, imaging acquisition parameters were kept constant for all conditions. Ciliary and cell membrane surface intensities were calculated in Image-J.

### Electrophysiological recordings

For a conventional whole-cell patch clamp experiment, tetracy-cline-inducible human embryonic kidney (HEK) 293 was used. The glass electrodes (Sutter instruments, BF150-86-10) were prepared using micropipette puller (Sutter instruments, SU-P1000). After preparation, the tip of the glass electrodes were polished using micro forge (Narishige, MF-830), resulting in the bath resistance, 4-6 MΩ. For ciliary excised single channel recordings, mouse inner medullary collecting duct 3 (mIMCD-3) cells were used. The glass electrodes (Sutter instruments, BF150-75-10) were fabricated by identical micropipette puller. The tip of the glass electrodes was forged until the bath resistance reached to 20-26 MΩ.

Data was acquired using an amplifier, Multiclamp 200B (Molecular Device, Axon Instruments, 200B), filtered at 5kHz. Data obtained using the amplifier was digitized at 10 kHz using Ditidata (Molecular Device, Axon Instuments, 1324A). The reference electrode was grounded using 3K KCl aga-bridge.

For the whole cell patch clamp, the step pulse protocol and the ramp pulse were applied to HEK 293 cells. The step pulse protocol consisted of voltage steps from −100 mV to +180 mV in 20 mV increments. The length of each step was 150 milliseconds. Holding potential and the tail pulse were given at 0 mV and −80 mV, respectively. The ramp pulse protocol was gradually applied from −100 mV to +180 mV with 0 mV holding potential during 500 milliseconds.

The intracellular solution contained (in mM): 90 Sodium Methanesufonate (NaMES), 10 Sodium Chloride (NaCl), 10 HEPES, 5 EGTA, 2 Magnesium Chloride (MgCl2) and 0.1 Free Calcium adjusted to pH 7.4 using NaOH and 290±5 mOsm/kg using D-mannitol. The extracellular solution contained (in mM): 145 mM Sodium Gluconate (NaCl), 5 mM Potssium Chloride (KCl), 2 mM Calcium Chloride (CaCl2), 1 mM Magnesium Chloride (MgCl2), 10 mM HEPES adjusted to pH 7.4 using NaOH and 290±5 mOsm/kg using D-mannitol. Osmolarity of the solutions were measured by vapor pressure osmometer (VAPRO, Wescor Inc.).

## Data analysis

The whole cell patch clamp and ciliary excised single channel data were recorded using Clampex 10.0 and analyzed using Clampfit 10.0 (Molecular Devices).

For the whole cell patch clamp, the current (I; pA) and voltage (V; mV) relationship was plotted with maximum current amplitudes obtained at each step pulse divided by capacitance (pF). For the comparison of the current density, maximum current amplitudes at +180 mV divided by capacitance (pF) were plotted and compared in the bar graph using Prism 5.0.

For the ciliary excised single channel recording, the current amplitudes were plotted in the conventional histogram and fitted to Gaussian equation at each potential.

After fitting the gaussian fitting, subtract µ1 from µ2 and plot the value on each voltage potential to show the current (I; pA) and voltage (V; mV) relationship. To calculate the conductance (G; pG), the currents plotted on voltage potentials were fitted to the linear equations. To calculate open probability of single channel activity, 10 seconds of the recording were selected on each voltage potential and calculate the open probability using the equation below:

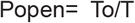

where To is the total time that the channel presented in the open state and T is the total observation time. If a patch contains more than one of the same type of channel, Popen was computed by:

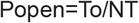

where, N indicates the number of channels in the patch. The following equation is used to populate data.

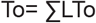

where, L indicates the level of the channel opening. The absolute probability of the channel being open NPo is computed by:

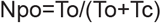

where, Tc indicates the total close.

## Computational Methods

The homomeric PC2 tetramer complex was simulated in presence of 7β,27-DHC oxysterols to sample free binding events, following up preliminary docking studies. The protein structure from PDB 5T4D3^5^ was embedded into 1-palmitoyl-2-oleoyl-glycero-3-phosphocholine (POPC) lipid bilayer and neutralized in 150mM of NaCl, using the CharmmGUI webserver72. Eight 7β,27-DHC molecules were placed in solution above the upper leaflet and below the lower leaflet. The total system size was ~300K atoms. Simulations were carried out using the ff14SB AMBER^73^ parameter set for the protein, the Joung–Cheatham^74^ parameters for the monovalent ions, LIPID17^75^ for the lipids, and the gaff^76^ force field for 7β,27-DHC. The TIP3P model was used to simulate the water. 7β,27-DHC geometry was first optimized, and its electrostatic potential was calculated with Gaussian v9-E016 with the 6–31 G* basis set. Next, the RESP charges were fitted into the electrostatic potential with antechamber and the force field parameters, topology, and starting coordinates were generated with AmberTools. Simulations were carried out with the pmemd MD engine on GPUs. Minimization consisted of 10,000 steps, switching from the steepest descent algorithm to conjugated gradient after 5000 steps. The system was then gradually heated from 0 to 303.15 K over 100 ps, and harmonic restraints with a spring constant of 10 kcal/mol/Å2 were applied to all heavy atoms except water oxygens. After reaching 100 K, we switched from NVT to NPT. After 100 ps, force constants of 10, 5, and 2.5 kcal/mol/Å2 were applied to the substrates and nearby residues (within 5 Å), protein, and lipid headgroups, respectively. Restraints were gently removed over the next ~10 ns, and the homomeric system was simulated for ~2 µs. Pressure (1 bar) was maintained using a semi-isotropic pressure tensor and the Monte Carlo barostat. Temperature was maintained with the Langevin thermostat with a friction coefficient of 1 ps−1. The SHAKE algorithm was used with a 2-fs time step. A non-bonded cutoff of 10 Åwas used, and electrostatics were calculated using the particle mesh Ewald method.

## Supplementary Information

**Extended Data Figure 1:** Ciliary oxysterol, 7β,27-DHC, only induce currents from sPC-1/2 overexpressing HEK293 cells

**Extended Data Figure 2:** 7β,27-DHC does not further potentiate the gain of function mutant, PC-2_F604P_

**Extended Data Figure 3:** Representative histogram collecting open channel events from the dose dependent experiment of figure 2.

## ACKNOWLEDGMENTS

This work was supported by National Institute of Health Grant R01DK127277 (MD and EC), National Research Foundation of Korea (NTF) grant funded by the Korean government (MSIT) (No.2019R1A6A3A03033302) (KH) and PKD Foundation (A137178) (KH). A portion of this work was performed under the auspices of the U.S. Department of Energy by Lawrence Livermore National Laboratory under Contract DE-AC52-07NA27344 (PB).

## Notes

### Competing Interest Statement

The authors have declared no competing interest.

